# Chromatin histone modifications and rigidity affect nuclear morphology independent of lamins

**DOI:** 10.1101/206367

**Authors:** Andrew D. Stephens, Patrick Z. Liu, Edward J. Banigan, Luay M. Almassalha, Vadim Backman, Stephen A. Adam, Robert D. Goldman, John F. Marko

**Affiliations:** Department of Molecular Biosciences, Northwestern University, Evanston, IL 60208; Department of Physics and Astronomy, Northwestern University, Evanston, IL 60208; Institute for Medical Engineering and Science, Massachusetts Institute of Technology, Cambridge, MA 02139; Department of Biomedical Engineering, Northwestern University, Evanston, IL 60208; Department of Cell and Molecular Biology, Northwestern University Feinberg School of Medicine, Chicago, IL 60611

## Abstract

Nuclear shape and architecture influence gene localization, mechanotransduction, transcription, and cell function. Abnormal nuclear morphology and protrusions termed “blebs” are diagnostic markers for many human afflictions including heart disease, aging, progeria, and cancer. Nuclear blebs are associated with both lamin and chromatin alterations. A number of prior studies suggest that lamins dictate nuclear morphology, but the contributions of altered chromatin compaction remain unclear. We show that chromatin histone modification state dictates nuclear rigidity, and modulating it is sufficient to both induce and suppress nuclear blebs. Treatment of mammalian cells with histone deacetylase inhibitors to increase euchromatin or histone methyltransferase inhibitors to decrease heterochromatin results in a softer nucleus and nuclear blebbing, without perturbing lamins. Oppositely, treatment with histone demethylase inhibitors increases heterochromatin and chromatin nuclear rigidity, which results in reduced nuclear blebbing in lamin B1 null nuclei. Notably, increased heterochromatin also rescues nuclear morphology in a model cell line for the accelerated aging disease Hutchinson-Gilford progeria syndrome caused by mutant lamin A, as well as cells from patients with the disease. Thus, chromatin histone modification state is a major determinant of nuclear blebbing and morphology via its contribution to nuclear rigidity.

## Introduction

The nucleus is an essential cellular structure that houses the genome and maintains its 3D structural organization, thus dictating gene transcription and cellular behavior. Disruption of genome organization and nuclear morphology occurs in many major human diseases including heart diseases, cancers, and laminopathies (Butin-Israeli et al., 2012; Reddy and Feinberg, 2013). Abnormal nuclear morphology has been used as one of the gold standards for cancer diagnoses for nearly a century in tests like the Pap smear (Papanicolaou and Traut, 1997). However, we still do not understand the underlying mechanical bases for these observed nuclear shape abnormalities. Most diseases that present abnormal nuclear morphology involve alterations of both major nuclear mechanical contributors, lamins and chromatin. Currently, it is widely held that nuclear disruptions are due to alterations of lamins, nuclear intermediate filaments that contribute to nuclear structure and mechanics (Shimi et al., 2008; Swift et al., 2013; Denais et al., 2016). It is also known that chromatin is an essential component of nuclear mechanical response (Chalut et al., 2012; Furusawa et al., 2015; Schreiner et al., 2015; Stephens et al., 2017; Banigan et al., 2017), which raises the question of whether chromatin’s contribution to nuclear mechanics affects nuclear morphology as well. It is therefore critical to identify the contributions of chromatin and disentangle them from those of lamins to understand the mechanical basis of aberrant nuclear morphology that is relevant to human disease.

Two major structural and mechanical components of the nucleus are the peripheral nuclear lamina and the interior chromatin. The nuclear lamina consists of four distinct lamin intermediate filament proteins, A, C, B1, and B2, that form separate but interacting networks at the periphery just under the inner nuclear membrane (Shimi et al., 2015; Turgay et al., 2017). Chromatin (DNA and associated proteins) fills the nucleus, with its organization dictated in part by attachments to the nuclear periphery/lamins (Guelen et al., 2008; Harr et al., 2015), crosslinking within itself (Denker and de Laat, 2016), and histone-modification-based compaction (Lleres et al., 2009; Stypula-Cyrus et al., 2013). Together, these two structural components determine separate mechanical response regimes of the nucleus to external forces. Chromatin controls the resistance to small deformations while lamin A dictates the nuclear strain stiffening that dominates resistance to large deformations (Stephens et al., 2017). While the separate force contributions have now been established, the roles of lamins and chromatin in maintaining nuclear structure remain unclear.

The most studied types of nuclear morphological abnormalities are protrusions from the normally ellipsoidal nucleus, termed ‘blebs’. To date, nuclear blebs have been mainly observed in cells with lamin perturbations, such as lamin A depletion (Sullivan et al., 1999), lamin B1 depletion (Lammerding et al., 2006), or lamin A mutations (Goldman et al., 2004). These blebs have been characterized as lacking lamin B1, exhibiting a distended lamin A network, and containing decompact euchromatin (Butin-Israeli et al., 2012). Currently, the prevailing hypothesis for the mechanism of bleb formation is that alterations to lamins destabilize the lamina, either passively (Funkhouser et al., 2013) or through response to mechanical perturbations (Wren et al., 2012; Cao et al., 2016). In the former model, phase separation of lamins A and B lead to aberrant structure, whereas in the latter, the less mechanically robust nuclear envelope is distended and/or ruptured by external forces. This latter mechanism of external-force-induced nuclear rupture and subsequent blebbing in nuclei with lamin perturbations has been shown to occur during cell migration through narrow channels (Denais et al., 2016; Raab et al., 2016) or via actin-based confinement in the cell (Le Berre et al., 2012; Tamiello et al., 2013; Hatch and Hetzer, 2016). However, to date there has not been an investigation of chromatin’s contribution to nuclear morphology, although altered histone modifications and chromatin decompaction are consistently present in both nuclei with abnormal morphology and nuclear blebs.

While previous studies have focused on lamin perturbations that induce blebs or abnormal morphology, these alterations also result in drastic changes to chromatin as well. Depletion of lamin B1 results in chromatin domain decondensation and loss of heterochromatin (Camps et al., 2014). Similarly, lamin A mutants associated with various laminopathies display loss of heterochromatin (Shumaker et al., 2006), disorganization of centromeres (Taimen et al., 2009), and/or increased chromatin fluctuations (Booth et al., 2015). In addition, nuclear sturdiness against rupture (Furusawa et al., 2015), nuclear morphological instability after chromatin digestion (Stephens et al., 2017; Banigan et al., 2017), and nuclear envelope dynamics (Schreiner et al., 2015) all rely on chromatin, its histone modifications, and/or its compaction state. Given that measurements in these reports demonstrate that chromatin directly contributes to nuclear mechanics, we hypothesized that chromatin state might also play a major causative role in nuclear bleb formation. Further support for this idea follows from recent work showing that disruption of histone linker H1 via overexpression of HMGN5 leads to bleb formation *in situ* and *in vivo* (Furusawa et al., 2015). These observations raise the intriguing possibility that changes in chromatin state might be able to drive nuclear morphological disruption, independent of perturbations to lamins.

To directly address the role of chromatin in nuclear morphology, we used inhibitors of the enzymes that modulate histone modification state to mimic chromatin alterations that are observed in human diseases. We used histone deacetylase inhibitors (HDACi) to increase histone acetylation, which corresponds predominantly to decompacted euchromatin (Lleres et al., 2009; Felisbino et al., 2014) and a histone methyltransferase inhibitor (HMTi) to decrease methylation, which corresponds predominantly to more compacted heterochromatin (Miranda et al., 2009; Stypula-Cyrus et al., 2013). These trends – increased decompacted euchromatin given stronger histone acetylation, and more compact heterochromatin given stronger methylation – only summarize changes in chromatin state resulting from changes in histone modifications. However, they do reflect the overall effects of histone modifications on chromatin contributions to nuclear mechanics (Stephens et al., 2017).

Chromatin decompaction treatments in mammalian cell lines resulted in decreased nuclear rigidity and formation of nuclear blebs without requiring alterations to the amount or organization of lamins. Blebs were depleted of chromatin, but enriched in euchromatin, relative to the nuclear body, and they were preferentially observed at the major axis poles. Strikingly, treatment with histone demethylase inhibitors, which increased heterochromatin levels and chromatin-based nuclear stiffness, partially rescued normal nuclear morphology by decreasing the number of blebbed nuclei occurring due to lamin B1 depletion. We recapitulated this finding in a lamin A mutant progeria model, as well as in cells from patients with progeria, which suggests that chromatin organization may in fact have a dominant role in determining nuclear morphology in some disease processes. Our findings suggest that chromatin histone modifications and their contribution to nuclear rigidity act as a major determinant of nuclear blebbing and morphology and may underlie mechanisms of nuclear morphological abnormalities seen in many human diseases.

## Results

### Increased euchromatin or decreased heterochromatin softens nuclei

To determine if the histone modification state of chromatin contributes to nuclear structure and morphology, we first established its contribution to nuclear mechanics. Previously, our lab developed a novel single nucleus isolation and micromanipulation assay to measure the whole-nucleus extensional spring constant (Stephens et al., 2017). We utilized MEF cells null for the intermediate filament vimentin (V-/-) because they facilitate reliable isolation of a single nucleus from a living cell in minutes and do not require drugs to depolymerize actin. Moreover, isolated MEF V-/- nuclei display similar force response to MEF wild-type nuclei (Stephens et al., 2017). Nuclei are isolated from single living cells via local lysis with 0.05% triton. Upon isolation, the nucleus is attached to two micropipettes, one “pull” pipette to extend the nucleus and one “force” pipette to measure force (nN) via deflection of the micropipette, which has a pre-measured spring constant. This provides a force-extension plot in which the slope of the line is the nuclear spring constant, which is 0.52 nN/µm for untreated MEF V-/- nuclei (Figure 1, A and B).

**Figure 1.**
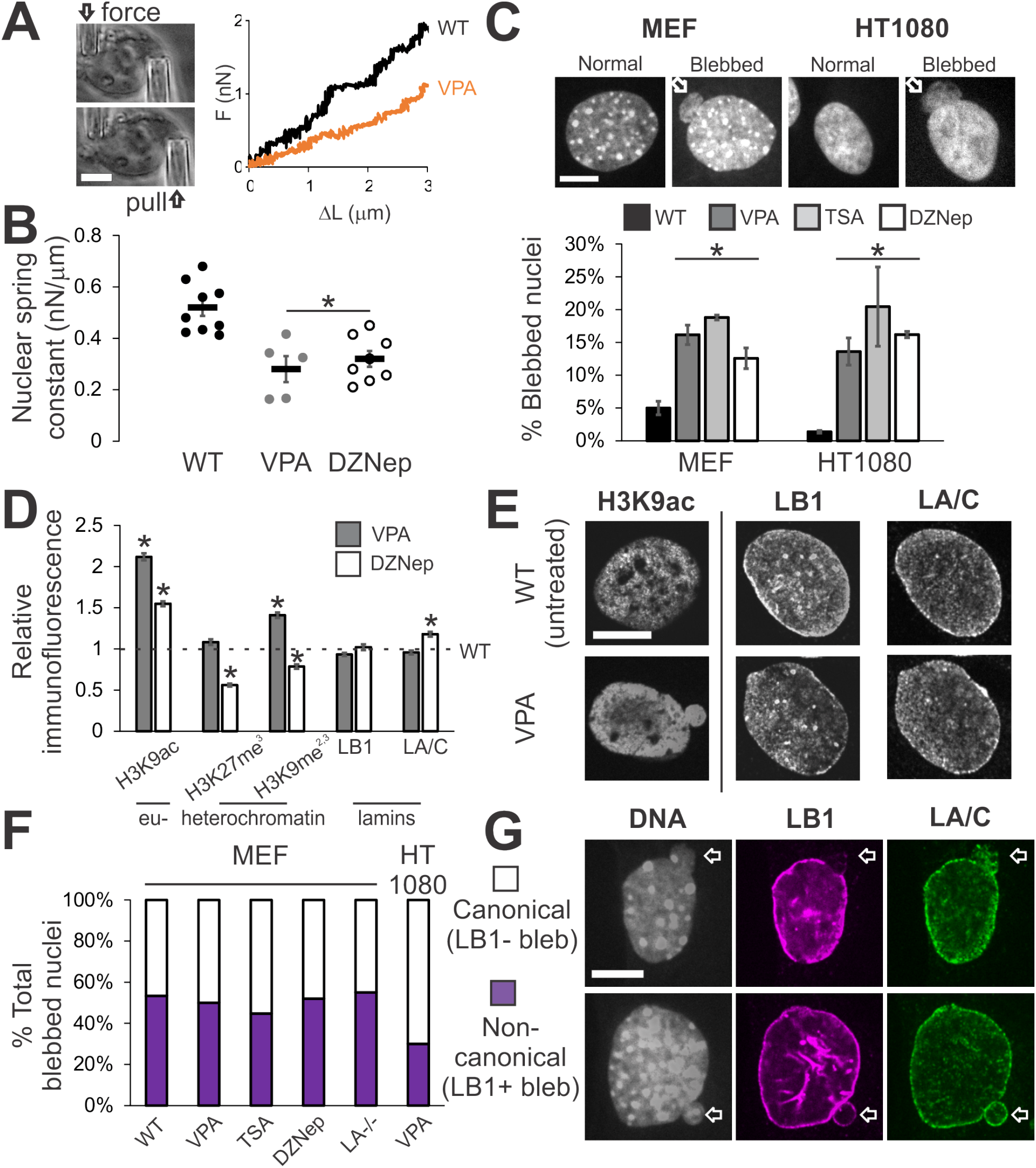
Increased euchromatin via HDACi or decreased heterochromatin via HMTi weakens nuclear rigidity and induces nuclear blebs, independent of changing lamin content and distribution. (A) Representative images of the micromanipulation force measurement technique and force-extension plot. Micromanipulation of a single isolated nucleus measures extension of the whole nucleus via movement of the “pull” pipette while simultaneously measuring force via the deflection of the “force” pipette with a known spring constant. (B) Nuclear spring constant, measured as the slope of the force-extension trace (nN/µm), for the chromatin-dominated initial force-extension regime (< 3 µm) for MEF V-/- nuclei untreated (WT, n = 9), increased euchromatin (VPA, n = 5), and decreased heterochromatin (DZNep, n = 8). (C) Representative images of normal and blebbed nuclei for MEF and HT1080 cells. White arrow denotes bleb. Percentages of blebbed nuclei in untreated (WT, black), increased euchromatin (VPA, gray and TSA, light gray), and decreased heterochromatin (DZNep, white; 3 data sets, total n > 400 for each condition) after 16-24 hours of treatment. (D) Relative intensities of euchromatin (H3K9ac), heterochromatin (H3K27me^3^ and H3K9me^2,3^), and lamins B1 and A/C measured via immunofluorescence relative to untreated (WT = 1 denoted by black dotted line), for VPA- and DZNep-treated MEFs (n > 90 for each). Western blots confirm immunofluorescence measurements (Supplemental Figure 4, A and B). (E) Representative immunofluorescence images of euchromatin (H3K9ac), lamin B1, and lamin A/C in untreated (WT) and VPA-treated MEFs. (F) Percentages of MEF wild-type (WT), VPA, TSA, DZNep, lamin A null (LA-/-), and HT1080 VPA-induced blebbed nuclei displaying canonical absence (white) or non-canonical presence (purple) of lamin B1 staining in the blebs (n = 15, 66, 38, 27, 20, and 30 respectively). (G) Representative MEF images of nuclei stained for DNA with Hoechst (gray) and lamin B1 (purple) and lamin A (green) via immunofluorescence. Examples of canonical blebs (top) lack lamin B1 and display distended lamin A, while non-canonical blebs (bottom) show no change in lamin B1 or A in the bleb. Scale bar = 10 µm. Error bars represent standard error. Asterisk denotes statistically significant difference (p < 0.01).

We then measured the nuclear spring constant in nuclei with decompacted chromatin. We used inhibitors of histone deacetylases and methyltransferases to alter histone modifications in ways that predominantly increase euchromatin or decrease heterochromatin (Strahl and Allis, 2000; Allis and Jenuwein, 2016). These treatments mimic changes in histone modification state observed in diseases displaying altered nuclear morphology (Butin-Israeli et al., 2012). Upon chromatin decompaction through increased euchromatin via treatment with the histone deacetylase inhibitor (HDACi) valproic acid (VPA, Figure 1D and Supplemental Figure 4A) (Lleres et al., 2009; Stypula-Cyrus et al., 2013; Felisbino et al., 2014), a significant decrease in the nuclear spring constant to 0.29 nN/µm was measured (Figure 1, A and B). Long-extension (> 3 µm strain) lamin-based rigidity via strain stiffening did not change upon VPA treatment (Supplemental Figure 1F). This finding is consistent with our previous experiments in which HDACi treatment decreases the chromatin-dominated nuclear spring constant for deformations smaller than 3 µm (Stephens et al., 2017).

Alternatively, chromatin decompaction can be achieved by decreasing the amount of compact heterochromatin via the broad histone methyltransferase inhibitor (HMTi) 3-Deazaneplanocin-A (DZNep). We refer to DZNep generally as decreasing heterochromatin because decreased histone pan-methylation primarily affects a number of heterochromatin markers (H3K9me^2,3^, H3K27me^3^, H3K79me^3^, and H4K20me^3^), with the notable exception of its effects on the active transcription marker H3K4me^3^ (Miranda et al., 2009; Black et al., 2012). To determine if loss of heterochromatin would also result in decreased chromatin-based nuclear stiffness, we performed micromanipulation force measurements on isolated nuclei from DZNep-treated cells. Similar to nuclei treated with HDACi, nuclei depleted of heterochromatin via DZNep (Figure 1D and Supplemental Figure 4B) exhibited a decreased nuclear spring constant of 0.31 nN/µm (Figure 1B). Similar to VPA treatment, DZNep treatment did not alter long-extension lamin-based rigidity (Supplemental Figure 1F). Thus, histone-modification-mediated chromatin decompaction via increased acetylation and/or decreased methylation governs the rigidity of the nucleus to short extensional stresses.

### Altered chromatin histone modification state and rigidity induces nuclear blebbing

Since perturbed nuclear mechanics are believed to induce blebbing, we tested whether decreased chromatin-based nuclear rigidity (Figure 1B) alone is sufficient to induce nuclear blebs. To do this, we assayed for nuclear blebs upon perturbation of histone modifications. Each nucleus was scored as blebbed if a protrusion larger than 1 µm in diameter was present (Figure 1C). Untreated wild-type MEF nuclei displayed blebs in 4% of nuclei, while the HT1080 human fibrosarcoma cell line displayed blebs in an even lower percentage of nuclei at 1% (Figure 1C, black bar).

Nuclei with alterations to histone modifications that result in decreased nuclear rigidity displayed an increase in nuclear blebbing. Cells treated with either HDACi valproic acid (VPA) or trichostatin A (TSA), displayed a significant increase in blebbed nuclei, with greater than 15% of nuclei displaying blebs in both MEF and HT1080 cell lines (Figure 1C, p < 0.001 χ^2^, n = 450-1000, gray bars). Similarly, the alternative approach of treatment with the broad HMTi DZNep resulted in a similar increase in the percentage of nuclei displaying blebs in both MEF and HT1080 cells to 12-18% (Figure 1C, p< 0.001 χ^2^, n = 200-600, white bar). MEF V-/- nuclei used to measure the nuclear spring constant display comparable nuclear blebbing percentages to MEF wild-type nuclei for both untreated and treated cells (Supplemental Figure 1A). While vimentin has been shown to enhance resistance to nuclear deformations and to maintain positional stability in cells (Neelam et al., 2015), we find that vimentin does not function to suppress blebs in MEF cells. Additionally, we note that other mammalian cell lines U2OS and HeLa showed a similar increased percentage of blebbed nuclei when treated with TSA (Supplemental Figure 1 B). This data suggests that decreasing the chromatin-based rigidity of the nucleus is sufficient to induce nuclear blebbing.

We surveyed common features of nuclear blebbing dynamics to better characterize nuclear blebbing via chromatin decompaction. Increased nuclear blebbing occurred as early as 2 hours after treatment, suggesting that cells do not require a round of division for the formation of nuclear blebs (Supplemental Figure 1C). Furthermore, live cell time-lapse imaging reveals nuclear rupture and bleb formation in G1 nuclei (Supplemental Figure 1D). Depolymerization of actin via cytochalasin D suppressed nuclear blebbing onset at 4-6 hours and significantly decreased nuclear blebs, from 15% to 4%, in cells treated with VPA for 24 hours (Supplemental Figure 1E). Lastly, average nuclear area and height did not change upon chromatin alterations, except for the area of DZNep-treated nuclei (Supplemental Figure 1, G and H). Thus, similar to the mechanism reported for nuclear bleb formation in lamin-perturbed cells, nuclear rupture accompanies bleb formation and actin confinement is required for blebbing (Vargas et al., 2012; Tamiello et al., 2013; Robijns et al., 2016; Hatch and Hetzer, 2016). Similar to these lamin perturbation studies, chromatin perturbations induce a comparable level of blebbing, with ~20% of cells displaying a blebbed nucleus. Altogether, these experiments show that chromatin-decompaction-induced nuclear blebs form in a similar manner to nuclear blebs induced by direct lamin perturbations.

### Chromatin alterations are sufficient to cause nuclear blebs independent of lamin perturbation

While it has been observed that lamin perturbations result in nuclear blebs, our experiments reveal increased nuclear blebbing can occur by directly altering chromatin instead of lamins. To assess overall changes to each nuclear component, we measured histone modification markers and lamin levels through immunofluorescence imaging. We performed immunofluorescence of a representative euchromatin marker H3K9ac and heterochromatin markers H3K27me^3^ (facultative) and H3K9me^2,3^ (constitutive). While immunofluorescence of the euchromatin marker H3K9ac increased as expected in HDACi-treated cells, both lamin B1 and lamin A/C staining were unchanged relative to wild-type (Figure 1, D and E, p > 0.05 t-test), consistent with previous findings (Stephens et al., 2017). Immunofluorescence experiments of MEFs treated with HMTi DZNep showed an expected decrease in the heterochromatin markers H3K27me^3^ and H3K9me^2,3^. These cells also exhibited a statistically significant increase in H3K9ac (euchromatin), but to a lesser degree than HDACi-treated cells (Figure 1D). Lamin B1 levels again remained unchanged, while lamin A/C increased by 20% in DZNep-treated cells. Western blots confirmed the immunofluorescence results (Supplemental Figure 4, A and B). These data show that decreasing lamin amounts is not necessary for the formation of nuclear blebs and that chromatin-state-based nuclear perturbations alone are sufficient to cause nuclear blebbing.

We next investigated whether local changes to the lamina are required to form nuclear blebs. Previously, nuclear blebs have been defined as protrusions lacking lamin B but retaining lamin A/C (Shimi et al., 2008). Thus, we assayed for the presence of lamin B1 and A/C in nuclear blebs induced by decreased chromatin-based nuclear rigidity (Figure 1, A and B). Interestingly, nuclear blebs induced by VPA treatment did not show loss or disruption of lamin B1 or A/C in 50% of MEF and 30% of HT1080 blebs (Figure 1, F and G). The presence of lamin B1 occurs with similar frequency (40-50%) in blebs observed in MEF wild-type, cells treated with TSA, cells treated with DZNep, and lamin A null cells (Figure 1F). This finding, combined with the lack of lamin-specific perturbations made in our experiments, contrasts with the prevailing hypothesis that nuclear blebs are reliant on lamin disruption. Instead, these results show that decreased nuclear rigidity via increased euchromatin formation and/or heterochromatin depletion is sufficient to cause nuclear blebbing, independent of the loss of lamin B1 and/or A/C in the nucleus or the bleb itself.

### Blebs are enriched in euchromatin

The composition of the chromatin within nuclear blebs may provide additional insights into the mechanism of blebbing by providing another point of comparison between blebs in chromatin and lamin perturbations. We compared fluorescence intensities of different stains in the nuclear bleb and the nuclear body to quantitatively determine the relative chromatin composition of the bleb. We included analysis of blebs formed via lamin perturbation, lamin B1 null MEFs (LB1-/-, (Shimi et al., 2015)), to compare to blebs formed due to altered histone modification state. Analysis of Hoechst intensity reveals that nuclear blebs are depleted of DNA relative to the nuclear body (bleb to body ratio, VPA 0.65 ± 0.04, DZNep 0.52 ± 0.07, and LB1-/- 0.55 ± 0.03, Figure 2A). Thus, the overall amount of chromatin is decreased within the bleb.

**Figure 2.**
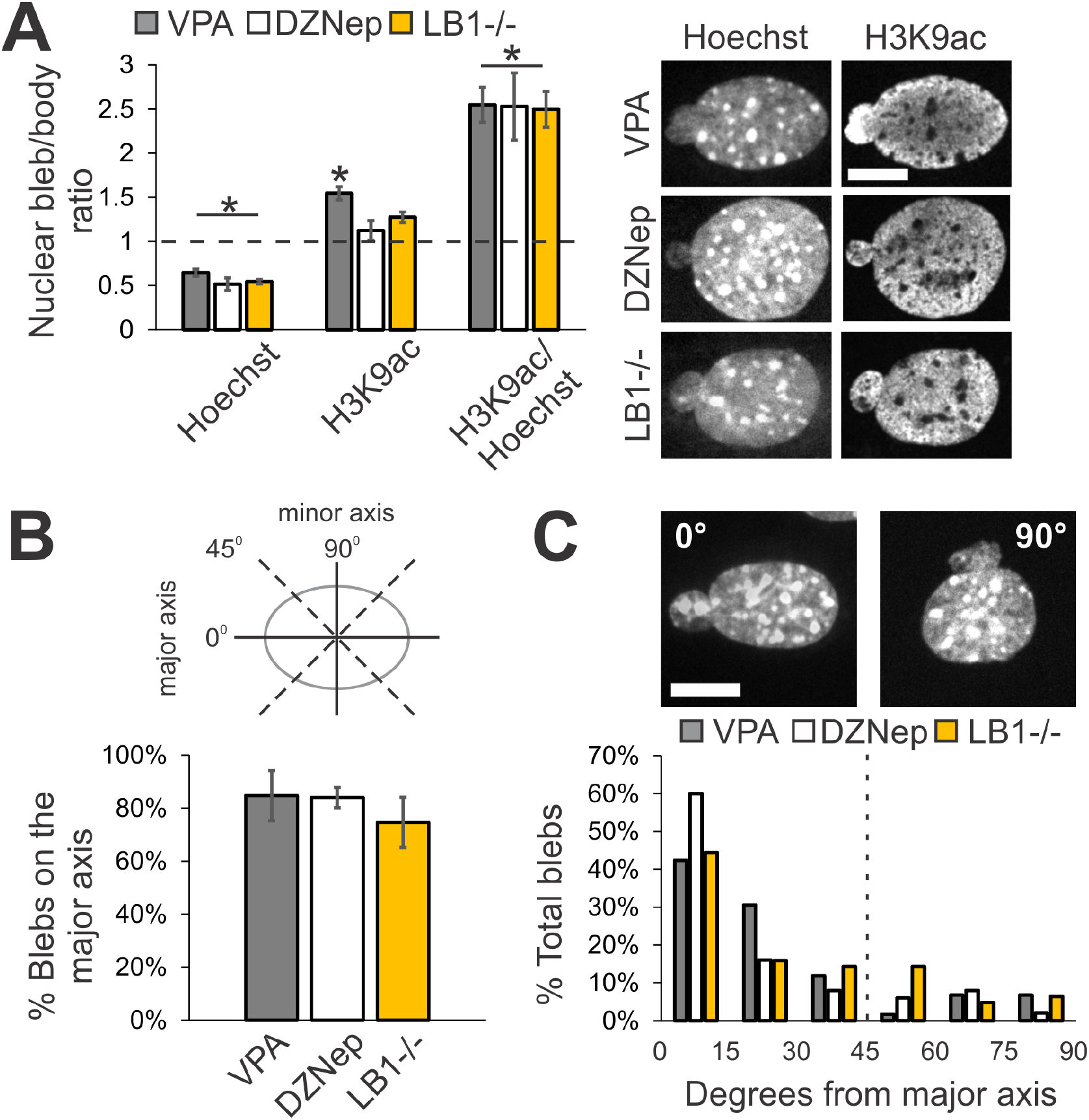
Nuclear blebs are enriched in euchromatin and form primarily along the major axis of the nucleus. (A) Fluorescence intensity ratio of nuclear bleb to nuclear body for Hoechst, H3K9ac, and H3K9ac divided by Hoechst (VPA n = 24, DZNep n = 10, and LB1-/- n = 22). Representative images are shown to the right. (B) Schematic of major vs. minor axis bleb location on the elliptical nucleus (average aspect ratio is 1.4). Average percentages of blebs that reside proximal to the major axis, within 45 degrees. (C) Representative images of blebs on (left) and off (right) the major axis of the nucleus. Histogram of bleb location on the nuclear body relative to the major axis, denoted as angle 0, for bins of 15 degrees (VPA n = 59, DZNep n = 50, and LB1-/- n = 63, two independent experiments each). Cells were treated with VPA or DZNep for 16-24 hours. Scale bar = 10 µm. Error bars represent standard error. Asterisk denotes statistically significant difference (p < 0.01, t-test).

Next, we investigated whether nuclear blebs are enriched in euchromatin, as has been reported for blebs arising from lamin-based perturbations (Dechat et al., 2008; Shimi et al., 2008). We measured the ratio of euchromatin marker H3K9ac signal in the bleb relative to that in the nuclear body. Nuclear blebs formed via VPA treatment show a statistically significant enrichment of H3K9ac in the bleb with a ratio of 1.54 ± 0.08 (VPA, n = 24, p < 0.001 t-test, Figure 2A). This increase could be due in part to the qualitatively observed enrichment of euchromatin at the nuclear periphery. Immunofluorescence imaging also reveals similar increased signal in the body and enrichment in the bleb of the euchromatin markers H3K9ac in HT1080 cells and H4K5ac in MEF cells treated with VPA (Supplemental Figure 2, A and B). Interestingly, in nuclei with decreased heterochromatin (DZNep or LB1-/-), which do not show H3K9ac enrichment at the periphery, nuclear blebs do not exhibit a statistically significant increase in H3K9ac in the bleb as compared to the nuclear body with ratios of 1.12 ± 0.11 and 1.27 ± 0.06, respectively (p > 0.05 t-test, n= 10 and 22, Figure 2A). However, it is nonetheless possible that other euchromatin marks are enriched within the bleb. More importantly, irrespective of enriched peripheral or homogenous H3K9ac spatial distribution, the relative ratio of euchromatin-to-DNA revealed an enrichment of ~2.5-fold within the bleb for all conditions (Figure 2A, H3K9ac/Hoechst). These results suggest that nuclear blebs are filled with decompact euchromatin irrespective of whether chromatin or lamins were directly perturbed.

### Blebs preferentially localize along the major axis

We hypothesized that if herniation of the nucleus were based on force balance then the most likely position of the bleb would be the region of greatest curvature and thus either the lowest local surface tension or highest local pressure (stress) – the major axis pole. To quantitate bleb position, we measured the angle of each bleb’s centroid relative to the major axis of its nuclear body, which was fit to an ellipse (average aspect ratio of normal nuclei is 1.4 ± 0.03; Figure 2B diagram). Histograms for nuclei of MEFs treated with VPA show that >40% of nuclear blebs are located within 15° of the major axis while 85% fall within 45° of the major axis (Figure 2, B and C, gray bar, n = 59, two experiments). Blebs that are positioned greater than 45° from the major axis are considered to be on the minor axis (Figure 2B). Bleb angular distribution was similar for DZNep-treated MEFs, with 60% located within 15° and 84% within 45° of the major axis of the nuclear body (Figure 2, B and C, white bar, n = 50, two experiments). Furthermore, VPA-treated HT1080 nuclei have a similar propensity for bleb location along the major axis (Supplemental Figure 2, A and C), providing preliminary evidence that this could be a general attribute of nuclear blebs. Even accounting for the increased angular weight at the pole due to the ellipsoidal shape of the nucleus does not explain the finding that blebs preferentially form on the major axis (Supplemental Figure 2, E and F). Blebs off axis did not show strong anti-correlation with nuclear body aspect ratio (Supplemental Figure 2D). We hypothesize that they appear due to fluctuations in nuclear shape or orientation that lead to transient off-axis positioning during imaging. Thus, chromatin-based nuclear blebs preferentially form along the major axis, the area of greatest curvature on the ellipsoidal nucleus.

We then assayed bleb position in MEF cells in which nuclear blebbing was induced by lamin B1 deletion (LB1-/-, (Shimi et al., 2015)) to determine if bleb positioning on the major axis is consistent across different types of nuclear perturbation (lamins vs. chromatin). In MEF LB1-/- blebbed nuclei, 75% of the blebs lie on the major axis, with an angular distribution similar to those of nuclei with histone modification state perturbations (Figure 2, B and C, yellow bar). This suggests that the physical mechanism for bleb formation and positioning may be consistent between blebs generated by direct perturbation to either chromatin or lamins. Blebs positioned along the major axis can also readily be seen in images from earlier papers reporting nuclear blebs, for example (Sullivan et al., 1999; Furusawa et al., 2015; Denais et al., 2016; Hatch and Hetzer, 2016), that did not quantitatively analyze blebs in this manner. Altogether, our data indicates that there is an underlying physical and/or biological mechanism that tends to localize blebs along the nuclear major axis.

### Lamin B1 depletion also decreases heterochromatin

Next, we analyzed chromatin histone modification state and blebbing in a lamin perturbation to determine if chromatin alterations are also present in previously observed blebbing scenarios. One of the most studied realizations of lamin-perturbation-based nuclear blebbing is lamin B1 depletion (Lammerding et al., 2006). While we have established that the presence of lamin perturbations is not required for bleb formation, it is possible that the eu/heterochromatin perturbations that we observe are present in all nuclear blebbing cases. We assayed heterochromatin amounts in MEF wild-type and lamin B1 null (LB1-/-) cells via immunofluorescence. Immunofluorescence signal from facultative heterochromatin marker H3K27me^3^ was drastically reduced by 80% in LB1-/- compared to wild-type, while the constitutive marker H3K9me^2,3^ did not significantly change (Figure 3A, n > 100 each). Our data are consistent with previous findings of decreased heterochromatin and decondensation of chromatin territories in lamin B1 depleted human cells (Camps et al., 2014). In summary, while MEF nuclei null for lamin B1 are missing a lamin component, these nuclei also have a drastic decrease in heterochromatin that, as shown by our experiments (Figure 1), is sufficient by itself to induce nuclear blebbing.

**Figure 3.**
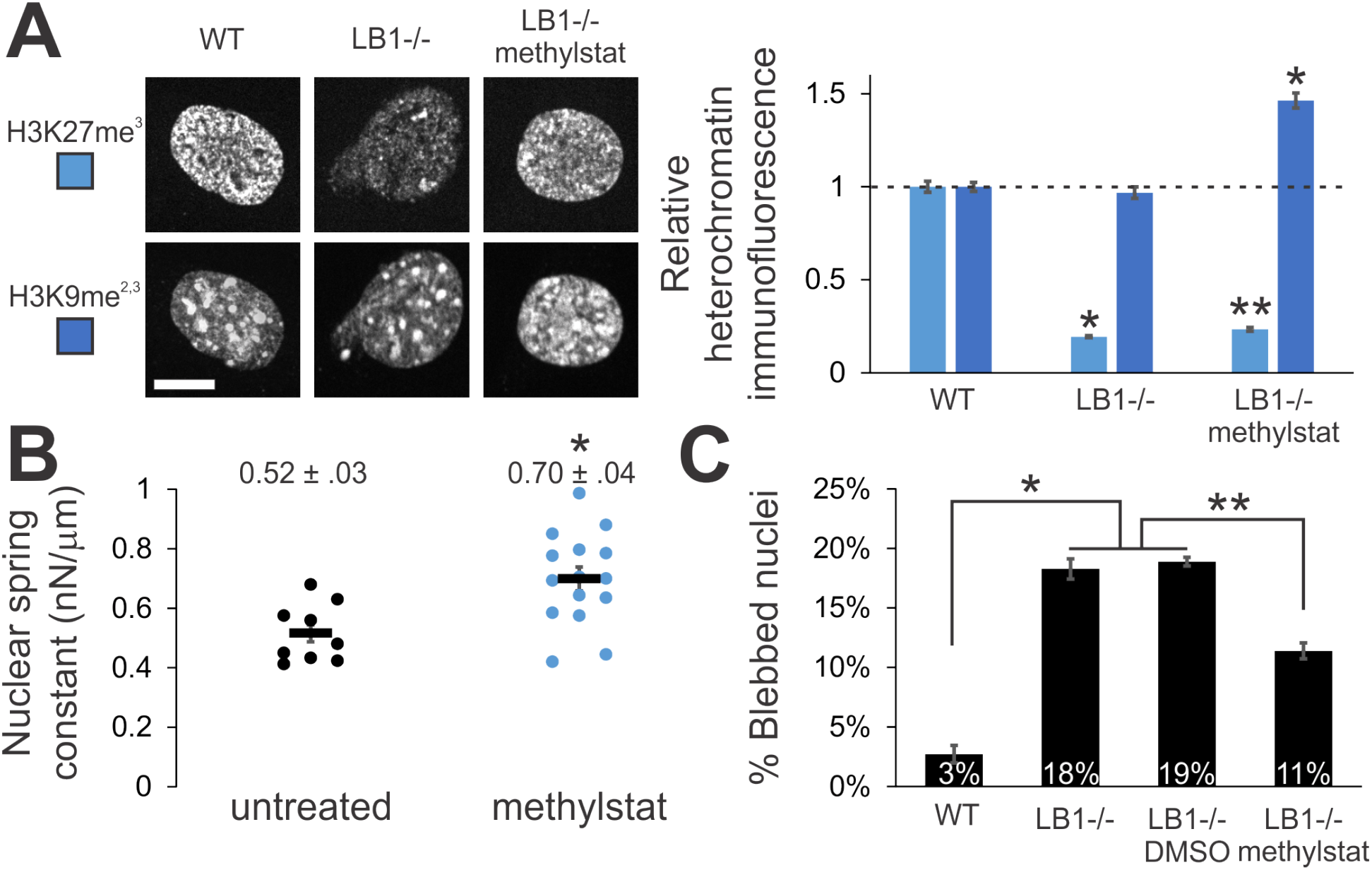
Increased heterochromatin via methylstat treatment increases nuclear rigidity and decreases nuclear blebbing in lamin B1 null nuclei. (A) Representative images of heterochromatin markers in MEF WT and LB1-/- nuclei untreated or treated with 1 µM methylstat for 48 hours. Plot to the right quantifies the change in relative fluorescence for H3K27me^3^ and H3K9me^2,3^ (n = 102. 106, and 112, respectively). Western blots confirm increased markers for heterochromatin while lamins remain unchanged for both MEF WT and LB1-/- treated with methylstat (Supplemental Figure 4, C and D). Addition of methylstat to cells in culture did not perturb cell growth (Supplemental Figure 3F). (B) Nuclear spring constants for MEF V-/- cells untreated and methylstat-treated via micropipette-based micromanipulation force measurements (n = 9 and 15, p < 0.005 t-test). Data for untreated cells is the same as in Figure 1. (C) Graph of percentage of nuclei displaying nuclear blebbing for MEF WT, LB1-/-, LB1-/- plus vehicle (DMSO), and LB1-/- treated with methylstat (n = 907, 901, 402, and 691, respectively). Scale bar = 10 µm. Error bars represent standard error. Asterisk denotes statistically significant difference (p < 0.01), and bars with different numbers of asterisks are also significantly different.

### Treatment with histone demethylase inhibitor methylstat increases heterochromatin formation, increases nuclear rigidity, and partially rescues morphology

Given that chromatin decompaction through altered histone modification state induces nuclear blebbing, we asked whether chromatin compaction via increased heterochromatin formation could rescue morphology. Methylstat is a broad histone demethylase inhibitor (HDMi) that primarily causes increased accumulation of heterochromatin markers (H3K9me^2,3^, H3K27me^3^, H3K79me^3^, H3K36me^3^, and H4K20me^3^), with the exception of its effects on the active transcription marker H3K4me^3^ (Luo et al., 2011). We treated MEF lamin B1 null (LB1-/-) cells with methylstat and measured the relative immunofluorescence of heterochromatin markers H3K27me^3^ and H3K9me^2,3^. Methylstat treatment resulted in an increase in both markers by 20% and 50%, respectively (Figure 3A). However, since LB1-/- nuclei have very little H3K27me^3^ signal, the 20% H3K27me^3^ increase upon methylstat treatment was small compared to the baseline wild-type level of H3K27me^3^. Nonetheless, the level of H3K9me^2,3^ in methylstat-treated LB1-/- cells was significantly higher than in wild-type cells (Figure 3A). Western blots of MEF WT and LB1-/- untreated and methylstat-treated cells confirm increased heterochromatin levels via these markers (Supplemental Figure 4, C and D). Thus, methylstat treatment is able to increase heterochromatin markers within nuclei null for lamin B1.

To determine if increased heterochromatin formation via methylstat would increase nuclear stiffness, we measured force response upon treatment in our standard MEF force measurement cell line (MEF V-/-). The short-extension nuclear spring constant increased upon methylstat treatment to 0.70 nN/µm from 0.52 nN/µm in untreated cells (Figure 3B, p < 0.01 t-test). This finding is consistent with our previous finding that increased heterochromatin (H3K27me^3^ and H3K9me^2,3^) stiffens HT29 nuclei (Stephens et al., 2017). Furthermore, long-extension rigidity attributed to lamins did not change upon methylstat treatment (Supplemental Figure 1F), similar to findings with VPA and DZNep. Thus, increased heterochromatin formation via methylstat treatment provides a method to increase chromatin-based short-extension nuclear rigidity.

Since heterochromatin formation increases nuclear stiffness, we investigated whether chromatin perturbations regulate nuclear morphology in nuclei with altered lamin content. To test this, we treated MEF LB1-/- cells with methylstat and assayed for nuclear blebs. MEF wild-type nuclei exhibited nuclear blebs in 3% of all nuclei while 18% of LB1-/- cells displayed blebs in their nuclei (Figure 3C, n > 400, 2-4 experiments each). Treatment with methylstat resulted in a significant decrease in the percentage of blebbed nuclei in MEF LB1-/- cells from 18% to 11% (p < 0.01, χ^2^), while treatment with the vehicle (DMSO) showed no change (Figure 3C, n > 400, 2-4 experiments each). Thus, increased chromatin-driven nuclear stiffness can rescue nuclear morphology, even in the presence of perturbed lamin content. Similarly, for LB1-/- nuclei treated with HDACi, which decreases nuclear rigidity, we measured an increase in nuclear blebbing from 19% to 31% (Supplemental Figure 3A, n > 600, p < 0.01, χ^2^), suggesting that chromatin-based alterations can both increase and decrease nuclear blebbing, even in a lamin perturbation. Taken together, these data suggest that lamin-based changes alone are not entirely responsible for nuclear morphology perturbations, or at least, they are not dominant over chromatin’s contribution to nuclear shape. Furthermore, these data show that chromatin histone modification state and its contribution to nuclear rigidity are major regulators of nuclear morphology.

### Increased heterochromatin via methylstat rescues nuclear morphology in a model of Hutchinson-Gilford progeria syndrome

Having established that chromatin-compaction-based nuclear rigidity driven by histone modifications provides a mechanism for nuclear morphology maintenance, we asked whether these observations could apply to human disease. A well-studied model of abnormal nuclear morphology is the accelerated aging disease, Hutchinson-Gilford progeria syndrome (HGPS), which is caused by mutations of lamin A. Overexpression of the most common HGPS mutant lamin A protein (LAΔ50), progerin, results in abnormally shaped nuclei (Goldman et al., 2004). Importantly, immunofluorescence reveals a drastic decrease of heterochromatin levels in nuclei expressing GFP-progerin. Relative to control GFP-lamin A expressing HeLa cells, GFP-progerin expressing HeLa cells displayed a loss of 60-70% in heterochromatin markers H3K27me^3^ and H3K9me^2,3^ via immunofluorescence (Figure 4, A and B, p < 0.01, t-test, n > 100) and Western blots (Supplemental Figure 4E), consistent with previous reports (Goldman et al., 2004; Shumaker et al., 2006). Thus, HeLa nuclei expressing GFP-progerin provide a clear case of abnormal nuclear morphology with perturbations to both lamins (mutant lamin A) and chromatin (decreased heterochromatin).

**Figure 4.**
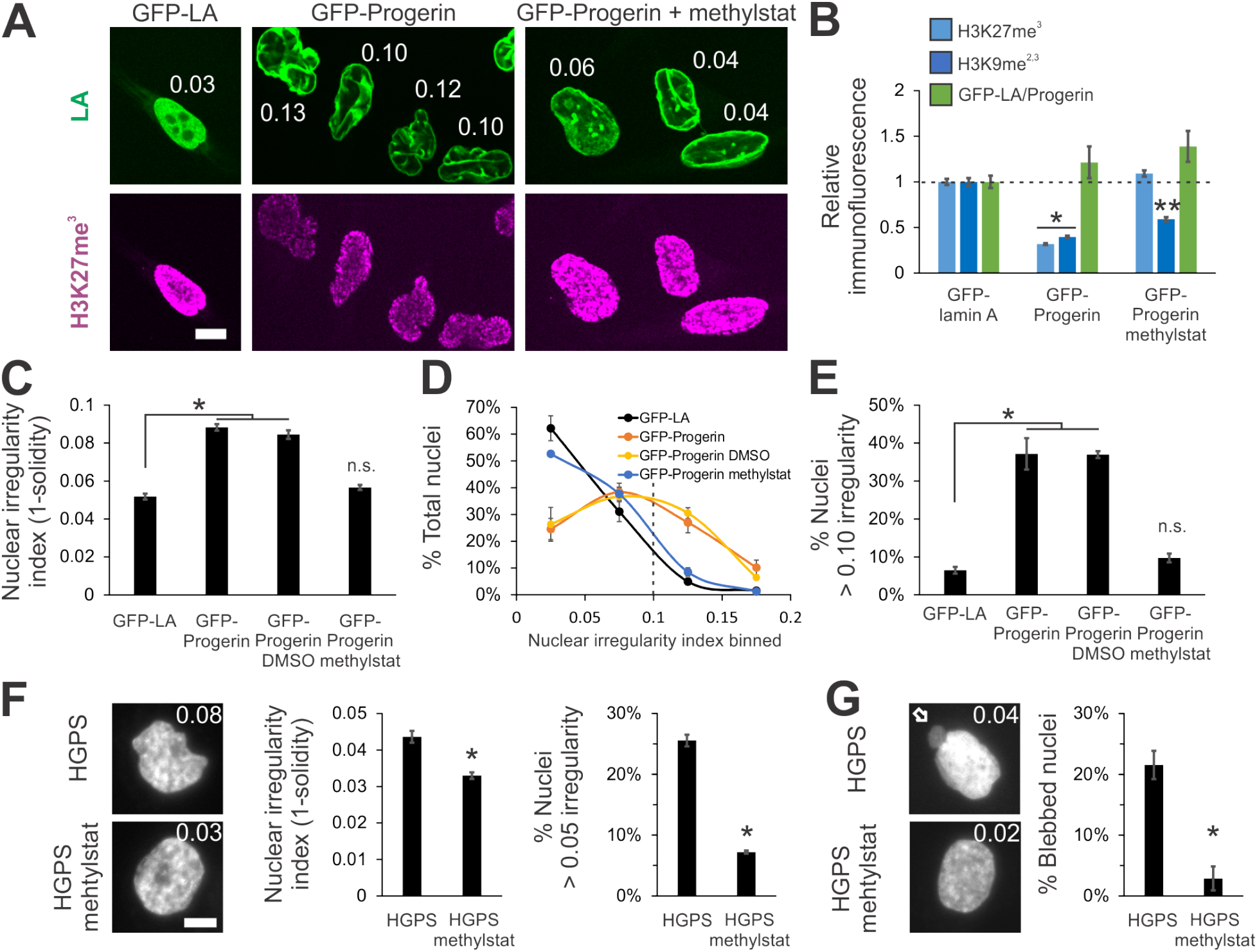
Increased heterochromatin via methylstat treatment rescues nuclear morphology in a model cell line and patient cells of laminopathy/aging disease Hutchinson-Gilford progeria syndrome caused by mutant lamin A. (A) Representative images of fluorescently tagged lamin A/progerin (green) and heterochromatin marker (H3K27me^3^, magenta) in HeLa nuclei expressing GFP-lamin A, GFP-progerin (GFP-LAΔ50), and GFP-progerin with methylstat for 48 hours. White numbers denote nuclear irregularity index of the corresponding nucleus (1 – *s*, where *s* = solidity = area/convex area). (B) Relative immunofluorescence for heterochromatin markers H3K27me^3^ and H3K9me^2,3^ (blue; n = 148 and 141, 135 and 146, and 126 and 104, respectively). (C, D, E) Plots of (C) average nuclear irregularity index, (D) histogram of nuclear irregularity index, and (E) average percentage of cells with irregularly shaped nuclei, which have irregularity index greater than 0.10 (HeLa GFP-LA n = 387, 2 data sets; HeLa GFP-progerin n = 656, 3 data sets; HeLa GFP-progerin DMSO n = 341, 2 data sets; HeLa GFP-progerin methylstat n = 496, 2 data sets). HeLa wild-type and HeLa GFP-LA nuclei have a similar average nuclear irregularity index (Supplemental Figure 3C). (F and G) Hutchinson-Gilford progeria syndrome (HGPS) passage 26 patient fibroblast cells untreated or treated with 2 µm methylstat for 48 hours and measured for (F) nuclear irregularity index and (G) nuclear blebbing (2 data sets, n = 130 and 139, respectively). Representative images have a white number denoting respective nuclear irregularity index and white arrow denotes nuclear bleb. Graphing nuclear irregularity index percentages > 0.05 was chosen because HGPS nuclei are more circular than HeLa nuclei. Scale bar = 10 µm. Error bars represent standard error. Asterisk denotes statistically significant difference (p < 0.01), where n.s. indicates no significant difference from control. Different numbers of asterisks are also significantly different.

The abnormal nuclear morphology due to expression of progerin perturbs shape across the entire nucleus instead of only discrete blebs. Therefore, we measured nuclear solidity (area divided by convex area), *s*, of Hoechst-labeled nuclei via live cell imaging to provide a measure of nuclear morphology. We then defined an irregularity index, 1 – *s*, to quantify the deviation of nuclear shape from a circle (the irregularity index of a circle is zero). HeLa control cells expressing GFP-lamin A have an average nuclear irregularity index of 0.052 (Figure 4, A and C, n = 387, 2 experiments, standard error approximately 0.003), which is similar to HeLa wild-type cells (Supplemental Figure 3C). As expected, HeLa cells expressing GFP-progerin have deformed nuclei with a higher average irregularity index of 0.088 (Figure 4, A and C, p < 1×10^−4^, t-test, n = 656, 3 experiments). To better quantify the amount of irregular nuclei, we calculated the percentage of nuclei with an irregularity index greater than 0.10 (denoted by dotted line in the histogram Figure 4, D and E). The percentage of irregular nuclei increased from 6 ± 1% in control HeLa cells to 38 ± 6% in those expressing GFP-progerin (Figure 4, D and E, p < 0.001, χ^2^).

To manipulate chromatin in these nuclei, we attempted to rescue heterochromatin levels in GFP-progerin expressing HeLa cells via the HDMi methylstat. We treated HeLa cells with methylstat coincident with active expression of GFP-progerin and then measured heterochromatin and GFP-progerin levels using immunofluorescence. Treatment with methylstat significantly increased the levels of both H3K27me^3^ and H3K9me^2,3^ heterochromatin markers. Immunofluorescence of H3K27me^3^ was rescued to levels similar to those measured in the control (GFP-LA, p > 0.05, t-test). Meanwhile, H3K9me^2,3^ signal in treated cells increased from 0.39 to 0.59 relative to control (Figure 4, A and B, p < 0.01, t-test, n > 100). The levels of GFP-progerin in the nucleus remained similar upon methylstat treatment (Figure 4, A and B, green, p > 0.05, t-test, n > 200). Western blots confirmed the immunofluorescence results (Supplemental Figure 4E). Thus, methylstat rescues heterochromatin levels, regaining much of the heterochromatin signal that was lost coincident with expression of GFP-progerin.

Next, we measured nuclear irregularity index in GFP-progerin expressing HeLa cells treated with methylstat to determine the contribution of chromatin to abnormal nuclear morphology in this disease model. Interestingly, methylstat and its associated rescue of heterochromatin levels also rescued nuclear morphology in HeLa cells expressing GFP-progerin, while the vehicle DMSO did not (Figure 4). In HeLa cells expressing mutant lamin A GFP-progerin and treated with methylstat, nuclear irregularity returned to control GFP-lamin A levels of 0.057 from 0.088 (Figure 4C, n = 496, two experiments, p > 0.01 t-test vs. GFP-LA). More strikingly, a histogram of irregularity index distribution shows a drastic difference between control GFP-lamin A and GFP-progerin nuclei, and a nearly full recovery of irregularity index across the entire population in GFP-progerin cells upon methylstat treatment (Figure 4D). Additionally, the percentage of irregular nuclei in GFP-progerin expressing cells was decreased to 7 ± 1% from 38 ± 6% upon restoration of heterochromatin levels by treatment with methylstat (Figure 4, D and E, p < 0.001, χ^2^). Thus, rescuing heterochromatin levels restores nuclear morphology in a model of the accelerated aging disease Hutchinson-Gilford progeria syndrome, which is driven by mutant lamin A. Taken together, these findings suggest chromatin is a major determinant of nuclear morphology that is relevant to human disease.

### Increased heterochromatin via methylstat rescues nuclear morphology in Hutchinson-Gilford progeria syndrome patient cells

To extend our findings, we investigated whether nuclear blebbing and morphology could be rescued in Hutchinson-Gilford progeria syndrome (HGPS) patient cells. Late passage (p 26) HGPS patient cells have previously been shown to present both abnormally shaped nuclei and nuclear blebs (Goldman et al., 2004) as well as a decrease in the heterochromatin marker H3K9me^3^ (Shumaker et al., 2006). We measured nuclear irregularity index to quantitate nuclear morphology in untreated and methylstat-treated cells. Similar to HeLa cells expressing GFP-progerin, treatment of HGPS patient cells with methylstat increased heterochromatin markers H3K9me^2,3^ and H3K27me^3^ (Supplemental Figure 4F) and rescued morphology, as measured by decreased nuclear irregularity index (Figure 4F). Abnormally shaped HGPS patient nuclei had an average nuclear irregularity index of 0.044 that decreased to 0.033 in methylstat-treated cells. This decrease is reflected by the decreased percentage of nuclei with a nuclear irregularity index above 0.05, which changed from 26 ± 3% to 7 ± 1% (Figure 4F, 2 data sets each, total n = 130 and 139, p < 0.001, χ^2^). Thus, increasing heterochromatin amounts using the broad HDMi methylstat rescues nuclear morphology in MEF lamin B null cells, in a HGPS model cell line, and in late-passage HGPS patient cells.

In addition to nuclear shape irregularity, HGPS patient nuclei also clearly present nuclear blebs. These blebs predominantly reside proximal to the major axis of the nucleus and display a decreased chromatin signal, as measured via Hoechst, of about half (0.52) compared to the nuclear body (Figure 4G, top image and Supplemental Figure 3, D and E), similar to other blebs (Figure 2). Increasing heterochromatin levels via methylstat treatment decreased nuclear blebbing, which demonstrates nuclear morphology rescue by a complementary measure in HGPS cells. Specifically, the percentage of cells with a nuclear bleb was drastically decreased from 22 ± 2% in untreated to 3 ± 2% in methylstat-treated cells (Figure 4G, 2 data sets each, total n = 130 and 139, p < 0.001, χ^2^). Thus, abnormal nuclear morphology and blebbing associated with a lamin A mutation in the laminopathy Hutchinson-Gilford progeria syndrome can be almost completely reversed by increasing the amount of compact heterochromatin within the nucleus. This finding, along with our other results, leads to our hypothesis that chromatin dictates nuclear morphology, even in situations in which lamins are also perturbed.

### Discussion

Nuclear morphology aids in the proper organization and maintenance of gene expression and is perturbed across a wide spectrum of human diseases (Butin-Israeli et al., 2012; Reddy and Feinberg, 2013). Chromatin and its histone modification state has been shown to determine nuclear mechanical response to small deformations (Stephens et al., 2017) and stability against buckling (Banigan et al., 2017). Consistent with this role, we show that chromatin histone modification state is a major factor determining the morphology of the nucleus, capable of generating euchromatin-filled blebs at the nuclear poles (Figures 1 and 2) or rescuing normal nuclear shape independent of lamin content or perturbation (Figures 3 and 4). These findings provide detailed evidence that chromatin is a major determinant of nuclear morphology relevant to both basic cell biology and human disease.

### Nuclear blebbing depends on chromatin and does not require lamin perturbation

Current models of nuclear blebbing do not account for the contribution of chromatin. Previous reports focusing on lamin perturbations suggested that lamin mechanics and the absence or disruption of lamins in blebs are the mechanical basis of bleb formation. For instance, although it is widely (though not universally (Raab et al., 2016)) held that blebs are defined by the lack of lamin B1 and the distention of the lamin A network (Butin-Israeli et al., 2012), this observation had not been rigorously quantified. Our assay for lamin B1 in blebs reveals that a significant portion of blebs, ~50%, in wild-type and chromatin-perturbed (HDACi- or HMTi-treated) nuclei contain both lamin B1 and A/C (Figure 1, F and G). Representative images from previous reports also reveal the presence of lamin B1 staining in blebs of nuclei depleted of chromatin remodeler BRG1 (Imbalzano et al., 2013) and nuclei of lamin A mutants (Taimen et al., 2009; Tamiello et al., 2013). Thus, nuclear blebbing does not require perturbation of lamins nor does it require the loss of lamin B within the bleb itself.

Instead, our experiments show that chromatin decompaction alone is sufficient to cause nuclear blebbing. Increasing euchromatin or decreasing heterochromatin decreases nuclear rigidity and increases nuclear blebbing in nuclei without necessarily altering the total or local lamin content (Figure 1). This is consistent with a previous study using HMGN5 overexpression to decompact chromatin that reported no change in bulk lamin levels, while nuclear blebs formed nonetheless (Furusawa et al., 2015). Similarly, this report found that nuclei with decompact chromatin were mechanically weaker, as measured by atomic force microscopy, and more likely to rupture during application of shear stress via syringe passes. Other studies show that alterations to chromatin remodelers, such as BRG1, cause abnormal nuclear morphology (Imbalzano et al., 2013), while chromatin compaction is critical to nuclear stability (Mazumder et al., 2008). These data all indicate that chromatin and its histone-mediated compaction are key contributors to the mechanics and morphology of the cell nucleus.

Furthermore, chromatin perturbations are consistently reported in cells exhibiting abnormal nuclear morphology, including in cases of lamin perturbations. These findings suggest that nuclear bleb formation in lamin-perturbed cells may be due to changes in chromatin compaction and dynamics (Camozzi et al., 2014), and may not be simply attributable to alterations in the lamina alone. For example, depletion of lamin B1 results in decreased heterochromatin (H3K27me^3^; Figure 3A) and decompaction of chromatin territories (Camps et al., 2014). Conversely, if lamin B1 loss in nuclear blebs is purely a symptom of chromatin-based decompaction and blebbing, then chromatin compaction changes may be driving nuclear bleb formation in diseases such as prostate cancer (Helfand et al., 2012).

Similarly, while lamin A loss does not directly alter the amount of heterochromatin (Pajerowski et al., 2007; Stephens et al., 2017), cells with mutations of lamin A show drastic changes in interactions between chromatin and the nuclear envelope (Dechat et al., 2008; Kubben et al., 2010; Kubben et al., 2012; McCord et al., 2013). Furthermore, laminopathies that display blebbing also show drastic changes in chromatin histone modifications and dynamics (Figure 4; (Goldman et al., 2004; Shumaker et al., 2006; Dechat et al., 2008; Booth et al., 2015)). Along with our experiments on progerin-expressing HeLa cells and HGPS patient cells (Figure 4), these observations suggest that morphological disruptions arising from lamin A perturbations may emerge due to the altered chromatin properties. Alterations to other nuclear proteins that alter morphology, such as WASH (Verboon et al., 2015), NAT10 (Larrieu et al., 2014; Robijns et al., 2016), BRG1 (Imbalzano et al., 2013), and nesprin (Kandert et al., 2007; Luke et al., 2008), also likely act in some capacity through their known effects on chromatin compaction. Thus, while perturbations to the lamina can induce nuclear blebbing, destabilization of nuclear shape may actually be due, at least in part, to lamin-induced disruption of chromatin interactions with the nuclear envelope.

### Nuclear blebbing is reversible

We have found that we can reverse or suppress the formation of nuclear blebs via chromatin-based nuclear rigidity driven by increased heterochromatin, the compact form of chromatin generated through histone modifications, largely due to the addition of methyl groups (Figures 3 and 4). This establishes that altered histone modification state and its contribution to nuclear rigidity can restore the typical shape of interphase nuclei. The rescue of lamin-based perturbations through compaction of the interphase chromatin adds to the data that blebs are dependent on the state of chromatin compaction. We hypothesize that the reversal of nuclear blebbing and abnormal morphology occurs through heterochromatin’s contribution to nuclear stiffness against external forces. Consistent with the idea of chromatin compaction rescuing nuclear morphology, chromatin condensation has been shown to occur during retraction of a bleb back into the nuclear body in a laminopathy mutant (Robijns et al., 2016). These findings support a possible pathway to rescue abnormal nuclear morphology, a major phenotype occurring in many human diseases, including progeria and cancer.

### Nuclear blebs occur in a subpopulation of cells

Nuclear blebbing occurring in both lamin and chromatin perturbations occur in a subset of nuclei, typically in the range of 10-25% of all nuclei. Specifically, MEF WT HDACi-treated nuclei displayed similar nuclear blebbing percentages to those of MEF lamin B null (LB1-/-) and HGPS patient cells at 20 ± 5% (Figure 1C, 3D, and 4G, respectively). This subpopulation size is consistent with many previous reports of blebbing percentages in lamin mutants (Lammerding et al., 2006; Vargas et al., 2012; Tamiello et al., 2013; Robijns et al., 2016; Hatch and Hetzer, 2016) and chromatin perturbations (Furusawa et al., 2015). Nuclear morphology as measured by nuclear irregularity also shows a distribution of shapes ranging from normal to abnormal for control HeLa cells, progerin-expressing HeLa cells, and HGPS patient cells (Figure 4, D and F). Interestingly, combining the bleb-causing perturbations of lamins and chromatin results in a significant increase in nuclear blebbing from 18% to 31% for LB1-/- treated with TSA (Supplemental Figure 3A) or HMGN5 overexpression in lamin mutants from 15% to 40% (Furusawa et al., 2015). This suggests that the subpopulation of blebbed cells can be expanded by multiple insults to nuclear stability.

Nonetheless, the relatively small size of the abnormal subpopulation may be due to the time and cell-cycle dependence and stochasticity of different mechanobiological processes. For example, stochasticity in rates of bleb formation and healing, as well as the steady-state balance between these two processes, could lead to apparently distinct normal and abnormal subpopulations. This stochasticity could be caused or enhanced by heterogeneity in cell behaviors such as migration, which leads to actin-based contractile stresses across the nucleus that vary over time and by cell. Other cellular processes that could influence the balance of subpopulations are nuclear envelope reformation post mitosis (Taimen et al., 2009; Samwer et al., 2017), cell age/passage (as in progeria (Goldman et al., 2004)), and transcriptional activity (Helfand et al., 2012). Future studies to investigate why blebs apparently form only in a small subpopulation of cells could provide further information about the mechanical and mechanistic processes driving bleb formation.

### Chromatin histone modification-based nuclear rigidity as a mechanism to maintain nuclear morphology

The rigidity of an object dictates its ability to maintain its shape. We find that this principle holds true for cell nuclei, which exhibit (or inhibit) abnormal shape, such as blebs, upon softening (or stiffening) due to chromatin alterations. These findings are consistent with our previous research showing that chromatin dictates nuclear force response to short deformations (< 3 μm, or about 30% strain) and prevents nuclear buckling, while lamin A mechanics are primarily important for resisting large deformations via strain stiffening (Stephens et al., 2017; Banigan et al., 2017). Here, we show that chromatin histone-modification-based nuclear rigidity regulates the ability of the nucleus to maintain its normal shape. Nuclei with high euchromatin or low heterochromatin levels are softer and succumb to blebbing, while nuclei with more heterochromatin are stiffer and resist blebbing (Figure 5).

**Figure 5.**
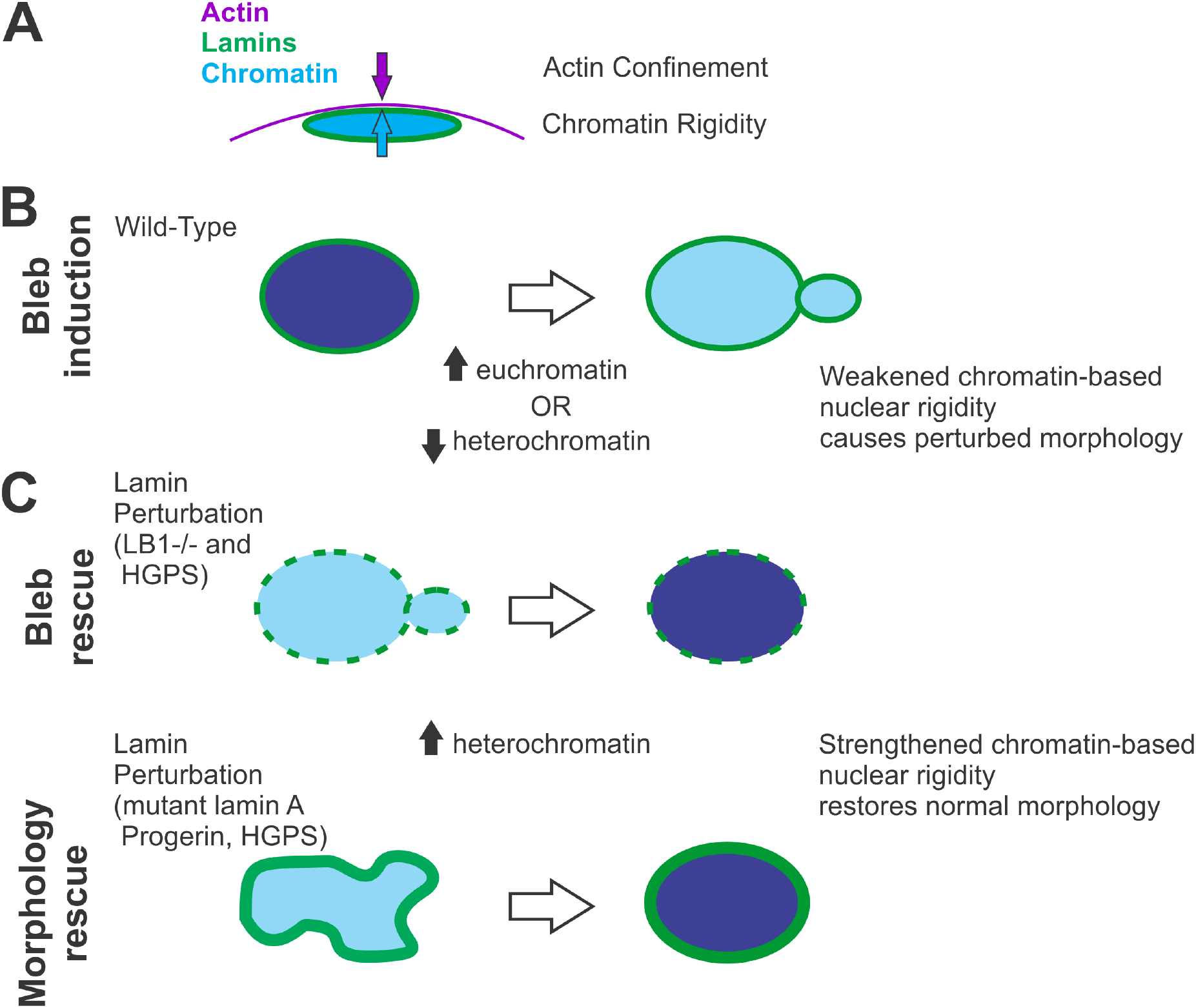
Alterations of chromatin-based nuclear rigidity affect nuclear blebbing and morphology, independent of lamin perturbation. (A) Side view depiction of force balance between actin confinement and chromatin-based nuclear rigidity. (B) Top view, weakening of the nuclear spring constant through chromatin decompaction via increased euchromatin or decreased heterochromatin causes the nucleus to herniate at the major axis pole. (C) Strengthening of the nuclear spring constant through chromatin compaction via increased heterochromatin partially rescues nuclear morphology observed in nuclei depleted of lamin B1. Similarly, chromatin compaction rescues nuclear morphology in nuclei overexpressing the mutant version of lamin A termed progerin that is associated with the accelerated aging disease Hutchison Gilford Progeria Syndrome (HGPS). Chromatin compaction through increased heterochromatin also rescues nuclear morphology by both decreasing nuclear blebs and abnormally shaped nuclei in HGPS patient cells.

Deforming forces for nuclei *in vivo* likely arise from extracellular compression or confinement (Le Berre et al., 2012) or from intracellular sources, such as the actin cytoskeleton (Supplemental Figure 1E; (Hatch and Hetzer, 2016)). For example, externally forced confinement of the nucleus occurs during cell migration as the cell squeezes through a narrow channel, which can lead to nuclear rupture and blebbing (Harada et al., 2014; Denais et al., 2016; Raab et al., 2016). Internally, actin cables running around and on top of the nucleus (Khatau et al., 2009) are believed to force nuclear confinement that is necessary to herniate the nucleus (Tamiello et al., 2013; Hatch and Hetzer, 2016) (Figure 5A, Supplemental Figure 1E). Furthermore, as demonstrated by cell spreading experiments, it is likely that the interplay of these external and internal factors contributes to the nuclear shape regulation (Li et al., 2015). In fact, cell spreading can amplify nuclear irregularities that arise in disease models (Tocco et al., 2017). Thus, altered nuclear rigidity should alter the force balance regulating nuclear morphology. This is consistent with our observation that nuclear blebs preferentially form at the major axis poles. This region of high curvature should have the highest local (outward) pressure and/or the lowest constraining surface tension (Figure 2, B and C). The novel observation in our experiments is that this disruption of force balance can be achieved via chromatin histone-modification-based alterations to nuclear rigidity (Figure 5, B and C).

Rigidity control through chromatin histone modification state provides a consistent mechanical explanation for nuclear blebs that cannot be understood through lamin mechanics alone. In particular, considering only lamin-based mechanics to understand nuclear morphology is potentially misleading. This is exemplified by the three main perturbations to the lamina that cause abnormal nuclear morphology, each of which has a different mechanical consequence. Depletion of lamins A or B1 or the mutation of lamin A all result in nuclear blebs or abnormal nuclear shape (Butin-Israeli et al., 2012). Lamin A is certainly a major contributor to nuclear mechanics, since when it is depleted nuclear stiffness is decreased, but its contribution is predominantly for large strains (> 30%) (Stephens et al. 2017). Lamin B1 loss, however, has been shown to have either no mechanical effect (Lammerding et al., 2006) or a slight stiffening effect on nuclei for large strains (Shin et al., 2013; Stephens et al., 2017). Complicating matters further, mutant lamin A progerin has been shown to stiffen nuclei (Dahl et al., 2006; Verstraeten et al., 2008; Booth et al., 2015). The lack of a clear connection between lamin mechanics and nuclear morphology is also illustrated by the fact that prelamin A expression in MEF lamin A null cells can rescue nuclear stiffness but cannot rescue nuclear morphology (Lammerding et al., 2006). Thus, current data does not support the idea that nuclear blebs arise from alterations of lamin-based mechanics alone.

Notably, abnormal nuclear morphology in the cases of lamin B1 loss and lamin A mutation occurs in conjunction with decreases in heterochromatin levels (Figures 3A and 4B, (Shumaker et al., 2006; Camps et al., 2014)). This heterochromatin decrease can lead to weakening of the nuclear spring constant in the chromatin-dominated regime (up to ~30% strain), which can account for the observed increase in frequency of nuclear morphological abnormalities (Figures 1 and 5B). Consistent with this picture, strengthening of chromatin-based mechanical response by increasing heterochromatin via methylstat rescues nuclear morphology in the same lamin perturbations (Figures 3 and 4, and Figure 5C). In summary, invoking changes to chromatin’s mechanical contribution to nuclear force response is sufficient to explain abnormal nuclear morphology arising due to lamin perturbations.

### Possible role of interactions between chromatin and the nuclear periphery in nuclear blebbing

While our data support a mechanism of nuclear morphology maintenance via chromatin-based nuclear rigidity (summarized in Figure 5), interactions between chromatin and the lamina/periphery also could affect or induce nuclear blebbing and abnormal morphology. Even though we have not directly explored the role of these interactions, loss of heterochromatin or generation of euchromatin in our experiments could result in a loss of connections between chromatin and the lamina (Dechat et al., 2008; Kubben et al., 2010; Kubben et al., 2012; McCord et al., 2013; Camozzi et al., 2014). This could inhibit chromatin-based nuclear rigidity, as observed in polymer Brownian dynamics simulations of nuclear mechanics (Stephens et al., 2017). This is consistent with our proposed rigidity loss mechanism for nuclear blebbing.

Alternatively, alterations to histone modification states could cause loss of chromatin-lamina connections that could promote deformations of the nuclear lamina itself. This idea parallels several experiments that disrupt heterochromatin and/or nuclear peripheral attachments (Kandert et al., 2007; Luke et al., 2008; Schreiner et al., 2015; Verboon et al., 2015). Consistent with this idea, experiments and simulations show that the lamina is susceptible to mechanical buckling when it is not coupled to a stiff chromatin structure (Banigan et al., 2017). Nuclear deformations due to loss of chromatin-lamina linkages could be further exacerbated by pressure gradients arising from external factors, such as confinement by the actin cytoskeleton or external environment (Raab et al., 2016; Cao et al., 2016). These open possibilities highlight the importance of better understanding interactions between chromatin and the nuclear periphery in future studies.

### Physiological implications of chromatin contribution to nuclear morphology

Cell nuclear mechanics and architecture are essential biophysical properties that influence cell behavior through their effects on transcription via mechanotransduction and regulation of gene spatial organization. We have shown that chromatin and its histone-modification-mediated compaction state are key factors underlying nuclear mechanics and morphology. The general observation that the histone modification profile and compaction state of chromatin are tightly controlled across several cell types suggests that cells also tightly regulate nuclear mechanics and shape. It remains to be determined whether altered rigidity, nuclear blebs, and other types of abnormal morphology drive disease or regulate normal functions, such as differentiation. Alternatively, these biophysical traits could be merely symptoms of cellular processes. Our results also suggest that lamin-based perturbations may alter nuclear morphology through their effects on chromatin, rather than through mechanical alterations to the lamina. Perhaps most intriguingly, the ability to regulate nuclear morphological hallmarks of disease via the compaction state of chromatin raises the possibility of new targets for therapeutic interventions.

## Materials and Methods

### Cell growth

MEF wild-type, MEF vimentin null (V-/-), HT1080, U2OS, and HeLa Kyoto cells were cultured in DMEM (Corning) complete with Pen Strep (Fisher) and 10% Fetal Bovine Serum (FBS, HyClone) at 37°C and 5% CO2. HGPS patient cells were cultured in MEM complete media with Pen Strep and 15% FBS (Goldman et al., 2004). HeLa progerin (GFP-LAΔ50) cells were grown in DMEM containing 1 mg/mL of G418 also at 37°C and 5% CO2. As outlined in (Taimen et al., 2009), progerin expression was induced via 2 *µ*g/mL doxycycline treatment for 48 hours. Upon reaching confluency, cells were trypsinized, replated, and diluted into fresh media.

### Histone modification drug treatment

Cells were treated with either histone deacetylase inhibitor valproic acid (VPA) at 1.5 µM or trichostatin A TSA at 100 nM for 16-24 hours to accumulate decompacted euchromatin, as previously reported in (Stephens et al., 2017). Cells were treated with histone methyltransferase inhibitor (HMTi) 3-Deazaneplanocin-A (DZNep) at 0.5 µM for 16-24 hours to deplete compact heterochromatin, as outlined in (Miranda et al., 2009). Cells were treated with histone demethylase inhibitor (HDMi) methylstat at 1 µM in MEF cells, 2.5 µM in HeLa cells expressing GFP-Progerin, and 2 µM Hutchinson-Gilford progeria syndrome patient cells for 48 hours, as generally outlined in (Luo et al., 2011). For validations, see immunofluorescence quantification in the main figures and Supplemental Figure 4 for Western blots. Treatment of MEF LB1-/- or HeLa GFP-progerin expressing cells with methylstat did not alter cell growth compared to untreated cells (Supplemental Figure 3, F and G).

### Micromanipulation force measurement of an isolated nucleus

Micromanipulation force measurements were conducted as described previously in Stephens et al. (Stephens et al., 2017). MEF vimentin null (V-/-) cells were used for their ease of nucleus isolation from a living cell and have similar nuclear force response as wild-type nuclei (Stephens et al., 2017). The nucleus was isolated by using small amounts of detergent (0.05% Triton X-100 in PBS) locally sprayed onto a living cell via a micropipette. This gentle lysis allows for a second micropipette to retrieve the nucleus from the cell via slight aspiration and non-specific adherence to the inside of the micropipette. Another micropipette was attached to the opposite end of the nucleus in a similar fashion. This “force” micropipette was pre-calibrated for its deflection spring constant, which is on the order of nN/µm. A custom computer program written in LabView was then run to move the “pull” micropipette and track the position of both the “pull” and “force” pipettes. The “pull” pipette was instructed to move 5 µm at 45 nm/sec. The program then tracked the distance between the pipettes to provide a measure of nucleus extension ~3 µm. Tracking the distance that the “force” pipette moved/deflected multiplied by the pre-measured spring constant provides a calculation of force exerted. Calculations were done in Excel (Microsoft) to produce a force extension plot from which the best-fit slope of the line would provide a spring constant of the nucleus (nN/µm). Each nucleus was stretched 2-4 times to provide an accurate and reproducible measurement of the nuclear spring constant.

### Immunofluorescence

Immunofluorescence experiments were conducted as described previously in Stephens et al. (Stephens et al., 2017). Cells were seeded on cover glasses in six well plates and incubated at 37°C and 5% CO2. Once at 80-90% confluence, cells were fixed with 4% paraformaldehyde (Electron Microscopy Sciences) in PBS for 15 minutes at room temperature. Cells were then washed three times for 10 minutes each with PBS, permeabilized with 0.1% Triton X-100 (US Biological) in PBS for 15 minutes, and washed with 0.06% Tween 20 (US Biological) in PBS for 5 minutes followed by two more washes in PBS for 5 minutes each at room temperature. Cells were then blocked for one hour at room temperature using a blocking solution consisting of 10% goat serum (Sigma Aldrich Inc) in PBS. Primary antibodies were diluted in the blocking solution at the following concentrations: H3K9ac 1:400 (C5B11, Cell Signaling), H3K9me2-3 1:100 (6F12, Cell Signaling), H3K27me^3^ 1:600 (C36B11, Cell Signaling), lamin A/C 1:10,000 (Active Motif), and lamin B1 1:500 (ab16048, Abcam). After being incubated with the primary antibodies overnight at 4°C in the dark, cells were washed with PBS three times for 5 minutes each. Next, cells were incubated with anti-mouse or anti-rabbit Alexa 488 or 594 (Life Technologies, 2 mg/mL) fluorescent secondary antibodies diluted at 1:500 in blocking solution for one hour at room temperature in the dark. Cells were washed with 1 µg/mL Hoechst 33342 (Life Technologies) in PBS for 5 minutes and then washed three more times with PBS. Finally, cover slides were mounted onto microscope slides using ProLong Gold antifade reagent (Life Technologies) and allowed to dry overnight at room temperature.

### Imaging and analysis

Immunofluorescence images were acquired with an IX-70 Olympus wide field microscope using a 60X oil 1.4 NA Olympus objective with an Andor iXon3 EMCCD camera using Metamorph. Image stacks with a step size of 0.4 *µ*m were also acquired using a Yokogawa CSU-X1 spinning disk Leica confocal microscope with a 63X oil 1.4 NA objective and a Photometrics Evolve 512 Delta camera using Metamorph. Exposure times for DAPI, Rhodamine and FITC were between 50-600 ms. Images were saved with Metamorph and transferred to ImageJ for analysis. Nuclei were selected by ImageJ threshold detection in the brightest plane or drawn by hand around Hoechst fluorescence if nuclei were too close together. Background fluorescence was determined by quantifying a 30×30 pixel area with no cells. Average intensity of the nucleus values were acquired and average background was subtracted using Excel. For comparisons between cell types or treatment conditions, relative intensities were reported as fold intensity relative to wild-type or untreated nuclei. For comparisons between nuclear blebs and nuclear bodies (Figure 2), the distinct nuclear portions were selected by hand drawing around Hoechst fluorescence. Following background fluorescence subtraction, the relative intensities of the bleb were reported as fold intensity relative to the nuclear body. A similar approach was taken to determine the presence or absence of lamin B1 in blebs, where loss of 50% of fluorescence was the threshold for distinguishing between the two classifications.

Bleb to body angle was calculated in ImageJ. First, the nuclear body was fit to an ellipse to provide the major axis and the angle at which the axis points. Next, a line connecting the centroid of the bleb to the centroid of the nuclear body provided the angle of the bleb. The angle of the bleb from the major axis was calculated as the difference between these two angles.

Statistical significance was determined for nuclear spring constants, immunofluorescence, and nuclear irregularity index measurements via the t-test. The chi-squared test was used to determine the statistical significance of changes in nuclear blebbing percentages.

### Live cell imaging

Cells were grown to the desired confluence in cell culture dishes containing glass coverslip bottoms (In Vitro Scientific). The dishes were treated with 1 µg/mL Hoechst 33342 (Life Technologies) for 10 minutes and then imaged on a wide field microscope as described above.

### Western blots

Western blots were carried out as described previously (Stephens et al., 2017). Protein was extracted via whole cell lysates (Sigma) or histone extraction kit (Abcam). Protein was loaded and run in 4-12 % gradient SDS-polyacrylamide protein gels (LICOR) for an hour at 100 V. Gels were then transferred to nitrocellulose blotting membrane with 0.2 µm pores (GE Healthcare) via wet transfer for 2 hours at 100 V. The membrane was then washed three times in TBST for 5 minutes each before blocking in either 5% Non-fat Milk or BSA in TBST for an hour at room temperature. Primary antibody was diluted into 5% Milk or BSA, depending on companies specifications, added to blotting membrane and allowed to shake and incubate overnight at 4⁰C. The same antibodies used in immunofluorescence experiments were also used for Western blotting (see above), except for anti-H3 (Cell Signaling Technology) and b-actin (LICOR). The next day the membranes were washed four times with TBST before incubation in secondary, antibodies conjugated to HRP (Millipore, 12-348 and 12-349), for 1 hour at room temperature. The membranes were again washed with TBST three times before chemiluminescence (PerkinElmer, Inc, NEL104001EA) and visualization using UVP imager. Quantification of Western blots was done in ImageJ.

## Acknowledgements

We thank Yixian Zheng for providing us with MEF LB1-/- cells (Shimi et al., 2015) and Aaron Straight for providing us with HeLa Kyoto cells. We thank Aykut Erbaş, Sumitabha Brahmachari, Haimei Chen, and Thomas O’Halloran for helpful discussions. A.D.S. is supported by NRSA postdoctoral fellowship F32GM112422 and was supported by a postdoctoral fellowship from the American Heart Association 14POST20490209. A.D.S., E.J.B., and J.F.M. are supported by NSF Grants DMR-1206868 and MCB-1022117, and by NIH Grants GM105847, CA193419, and via subcontract DK107980. S.A.A. and R.D.G. are supported by NIH GM106023, GM0969 and Progeria Research Foundation PRF 2013-51. L.M.A. and V.B. are supported by NIH grants R01CA200064, R01CA155284, NSF grant CBET-1240416, and Lungevity Foundation. This work was funded by the Chicago Biomedical Consortium with support from the Searle Funds at the Chicago Community Trust.

